# Nutrition induced changes in the microbiota can cause dysbiosis and disease development

**DOI:** 10.1101/2024.07.29.605727

**Authors:** Tim Lachnit, Laura Ulrich, Fiete M. Willmer, Tim Hasenbein, Leon X. Steiner, Maria Wolters, Eva M. Herbst, Peter Deines

## Abstract

The increasing global prevalence of inflammatory and autoimmune diseases highlights the need to understand their origins. The overfeeding hypothesis suggests that disturbances in the microbiota due to diet may initiate these diseases. Using the model organism *Hydra*, we established a causal link between environmental alterations in the microbiota and disease development. Relocating Hydra to natural lakes caused significant microbial shifts due to new colonizers and nutrients. Nutrient manipulation removed the competitive advantage of the well-adapted resident microbiota, disrupted its nutrient-blocking capacity and triggered specific microbiota changes leading to disease. L-arginine supplementation alone transformed *Pseudomonas* from a commensal microbe into a pathogen, showing pathogenicity as context-dependent. Our findings support the overfeeding hypothesis, highlighting the role of microbial and nutrient dynamics in disease development.

## Introduction

The prevalence of inflammatory and autoimmune diseases is increasing rapidly worldwide. Although therapeutic progress has been made, our understanding of the underlying causes remains limited. Another perspective that advances our understanding of the origin of these complex diseases is to consider that all eukaryotic organisms, including plants, animals and humans, are associated with complex microbial communities(*1*, *2*). Alterations in host-associated microbial communities are thought to lead to dysbiosis and disease development. For instance, several human diseases, including inflammatory bowel disease(*3*), multiple sclerosis(*4*), atopic dermatitis(*5*) and type 2 diabetes(*6*), are associated with alterations in microbial community composition. While a causal link between changes in the microbiota and disease aetiology has not yet been established, genetic predisposition is another factor that contributes to the development and manifestation of complex diseases(*7–9*). However, inherited genetic risk factors for complex diseases can only explain a small percentage of disease development, while environmental factors are considered to be the main trigger for disease onset(*10*). It is thought that the post-modern industrial lifestyle, with its excessive nutrient intake and high-calorie, processed foods, decouples natural host-microbe associations. Such changes result in functional alterations, increased growth of microbes and their byproducts, shifts in microbial community composition and ultimately facilitate the development of disease, as proposed by the “overfeeding hypothesis”(*11*). In addition, the loss of important microbes through antibiotics and the limited availability of new bacterial colonisers due to increased sanitation, as outlined in the “hygiene hypothesis”, may negatively impact the microbiota’s resilience and the acclimation potential of eukaryotic organisms to environmental changes. Our objective is to gain an understanding of how these environmental factors disrupt the homeostatic relationship between the host and its microbiota. This is of clinical relevance, as it is necessary to identify the biological principles that underpin the transition of commensal microbes to pathogens. Here, we experimentally investigate the intricate connection between environment-induced microbiota changes, nutrient conditions and disease development, using the freshwater polyp *Hydra* as a model organism. We used a multifaceted approach, combining observational field studies with controlled laboratory experiments, to reveal how ecosystems generate selective pressures that result in the emergence of a pathobiont (organism that can cause harm under certain conditions). In order to uncover the ecological drivers that lead to the selection of pathogenic traits, we also conducted nutrient manipulation experiments on both wild-type and mono-colonised organisms to establish causality and explore underlying mechanisms.

## Results

### The environment has a strong impact on microbial community composition

The present study aimed to investigate the effect of the environment on host-associated microbial communities. The freshwater polyp *Hydra vulgaris* was selected as a model organism due to its suitability for clonal line growth and the fact that the majority of its microbes reside on the outer mucosal surface of the organism, in direct contact and exchange with the environment(*12*). This makes it possible to track interactions of new colonisers with the host and to understand the influence of external nutrient supply on surface-associated microbes. The microbial community of *Hydra* has been subject to extensive study, with a well-documented host-specific composition(*13*, *14*). This community is regulated by the host immune system(*13*, *15–17*). The aim of the study was to investigate the ability of *Hydra* to maintain a host-specific microbial community composition under different environmental conditions. To achieve this, laboratory-grown polyps were transferred to lakes with different nutrient levels in the field (Fig. S1) and to lake water under controlled laboratory conditions (Fig. 1, A), with the microbial community composition of the polyps being monitored over time.

**Fig. 1.**
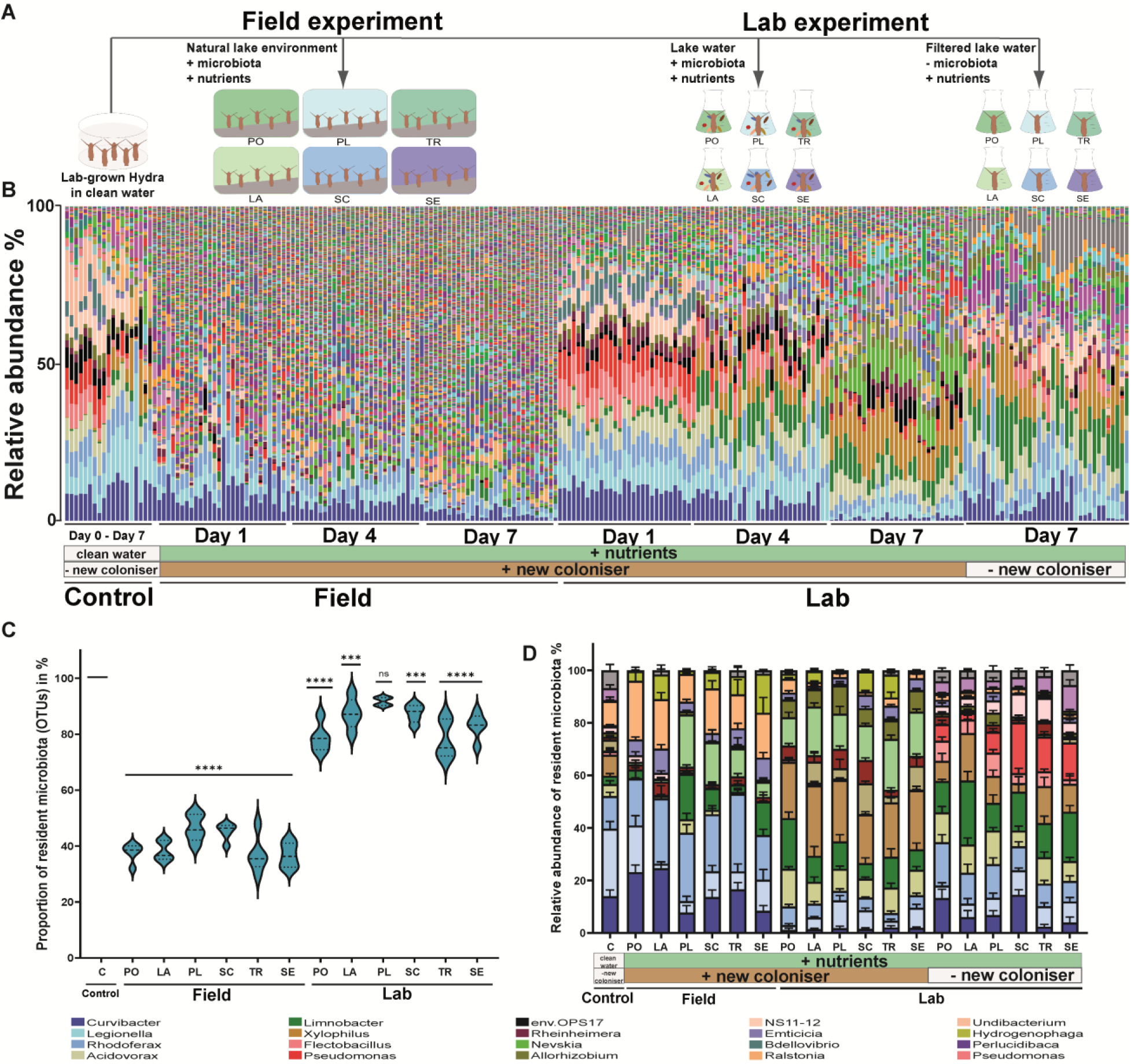
: Natural lake environments have a strong impact on *Hydra*’s microbial community composition. (A) Graphical illustration of the experimental design. Laboratory-grown *Hydra vulgaris* (AEP) were transferred to different lake environments in the field or to the corresponding lake water under laboratory conditions. In addition, laboratory-grown *Hydra* were exposed to sterile, filtered lake water in the absence of new microbes. (B) Bar-chart illustrating microbial community composition based on operational taxonomic units (OTUs). (C) Proportion of resident bacterial OTUs compared to new colonisers. Dotted lines in violin plots represent the median. One-Way ANOVA with Dunnett’s multiple comparison test compares each group with the control [P > 0.01 = not significant (ns); ***P < 0.001; ****P < 0.0001]. (D) Environmental changes affect the resident core microbial communities. Resident bacterial OTUs undergo abundance shifts in the field and when exposed to lake water in the laboratory. Exposing *Hydra* only to sterile, filtered lake water in the absence of any microbes already had an impact on microbial community composition. Data are mean ± s.e.

Exposure of laboratory-grown *Hydra* to different nutrient-rich lake water environments in the field and under controlled laboratory conditions for up to 7 days had a significant effect on the composition of the host-associated microbial community (Fig. S2). 16S rRNA gene sequencing analysis revealed that the associated microbial community of *Hydra* had already undergone a notable change in polyps exposed to lake water under laboratory conditions. This effect was even more pronounced in *Hydra* individuals that were exposed to natural field conditions (Fig. 1, B). The analysis was focused on bacteria that had been previously identified within the *Hydra* microbiota prior to lake environment exposure. Significant shifts were observed within the composition of this resident microbiota. The relative abundance of *Hydra*-specific bacteria decreased gradually over time. Starting from an initial abundance of 100 %, the total relative abundance under laboratory culture conditions declined to approximately 80 % after 7 days. In the field, the relative abundance decreased to approximately 40 % after 7 days (Fig. 1, C). Consequently, the relative abundance of resident microbes was higher when polyps were exposed to lake water under laboratory conditions. These changes in relative abundance were not solely caused by additional colonisation of bacteria from the surrounding lake water. By removing all new microbial colonisers from the analysis and setting the relative abundance of the resident microbiota to 100 %, it was demonstrated that the resident microbiota underwent compositional shifts in the lake environment (Fig. 1, D). Some bacteria were reduced in abundance (*Curvibacter* and *Legionella*), while others fluctuated in their relative abundance (e.g. *Rhodoferax*). In contrast, some microbes that were normally underrepresented under laboratory culture conditions (e.g. *Limnobacter* and *Nevskia*), increased in abundance manifold (Fig. 1, D).

Our observations indicate that new microbes can colonise and that the composition of the resident microbiota undergoes significant shifts, when exposed to different lake environments. This challenges the view of a stable host-specific microbial community, which has been thought to be regulated and maintained by host immunity(*13*, *18*, *19*). Similar findings stating that the environment is the primary determinant of the human microbiota composition have been reported by Gacesa *et al*.(*10*). Furthermore, it was demonstrated that the wider environment exerts a profound influence on an organism’s microbial community(*20*). Monitoring ecosystem-wide microbial trafficking underscores the significance of interactions between the environment and host microbial diversity for a holistic view on organism health.

### The chemical environment induces shifts in the resident microbiota

We hypothesised that host-associated bacteria are dependent on nutrients provided by their host, representing an important regulator of microbial growth in the mucosal environment(*11*). Additional nutrients from the environment can disrupt this microbe-host dependency, allowing for an uncontrolled growth of microbes, including those not specifically adapted to the host(*11*). Under laboratory conditions, *Hydra* is cultured in nutrient-deficient water(*21*), that lacks the natural nutrient sources for bacteria typically found in lake environments, such as dissolved organic matter (DOM) and particulate organic matter (POM). By removing microbes through filtration and exposing *Hydra* solely to the water chemistry of different lake environments, we were able to observe the influence of lake water nutrients on the resident microbiota, compared to a control group that lacked any additional nutrients.

The results demonstrate, that resident microbiota’s composition was already influenced by the water chemistry of different lake environments, even in the absence of new microbial colonisers (Fig. 1, D). In particular, the increase of nutrients in the water had a significant impact on the composition of the microbial community. More specifically, we observed a decrease in *Curvibacter* and *Legionalla*, while *Acidovorax*, *Xylophilus*, *Limnobacter* and *Pseudomonas* showed an increase in abundance compared to the control in nutrient-deficient water (Fig. 1, D). It is noteworthy that our findings revealed not only differences between lake water chemistry and nutrient-deficient water, but also variations among different lake waters in terms of microbial community composition. This variation can partly be explained by differences in dissolved organic carbon (DOC) and phosphate concentrations. By analysing the nutrient load and composition of the different lake waters, including measurements of DOC, total phosphate and total nitrogen (Table S3), we found that, for example *Acidovorax* and *Xylophilus*, exhibited a positive correlation with increasing DOC levels (Fig. S3), whereas *Rhodoferax* and *Flectobacillus* demonstrated a positive correlation with the total amount of phosphate (Fig. S4). Furthermore, it was observed that not only quantity but also composition of nutrients exerted a selective influence on the microbiota’s composition. The ratio of DOC to nitrogen or phosphate was calculated, which revealed that decreasing C/N ratios were positively correlated with the relative abundance of *Pseudomonas* and decreasing C/P ratios were positively correlated with *Curvibacter* (Fig. S5). Taken together these results suggest that nutrient load and composition have a selective effect on the composition of the host-associated microbiota.

### Elevation of nutrient levels induces dysbiosis and disease development

Additionally, we investigated the impact of nutrients on the host-associated microbiota. We exposed laboratory-grown *Hydra* to distinct nutrient regimes in the absence of new microbial colonisers. The artificial enrichment of nutrients in *Hydra* water (Fig.2, A) resulted in a change in the microbial community composition (Fig. S6). In both nutrient-enriched environments, the relative abundance of *Curvibacter* significantly decreased dropping from 60-80 % (control) to less than 7 % (Fig. 2, B). Concurrent with the loss of *Curvibacter,* an increase was observed in previously underrepresented *Pseudomonas, Duganella* and *Rheinheimera* (Fig. 2, B).

**Fig. 2:**
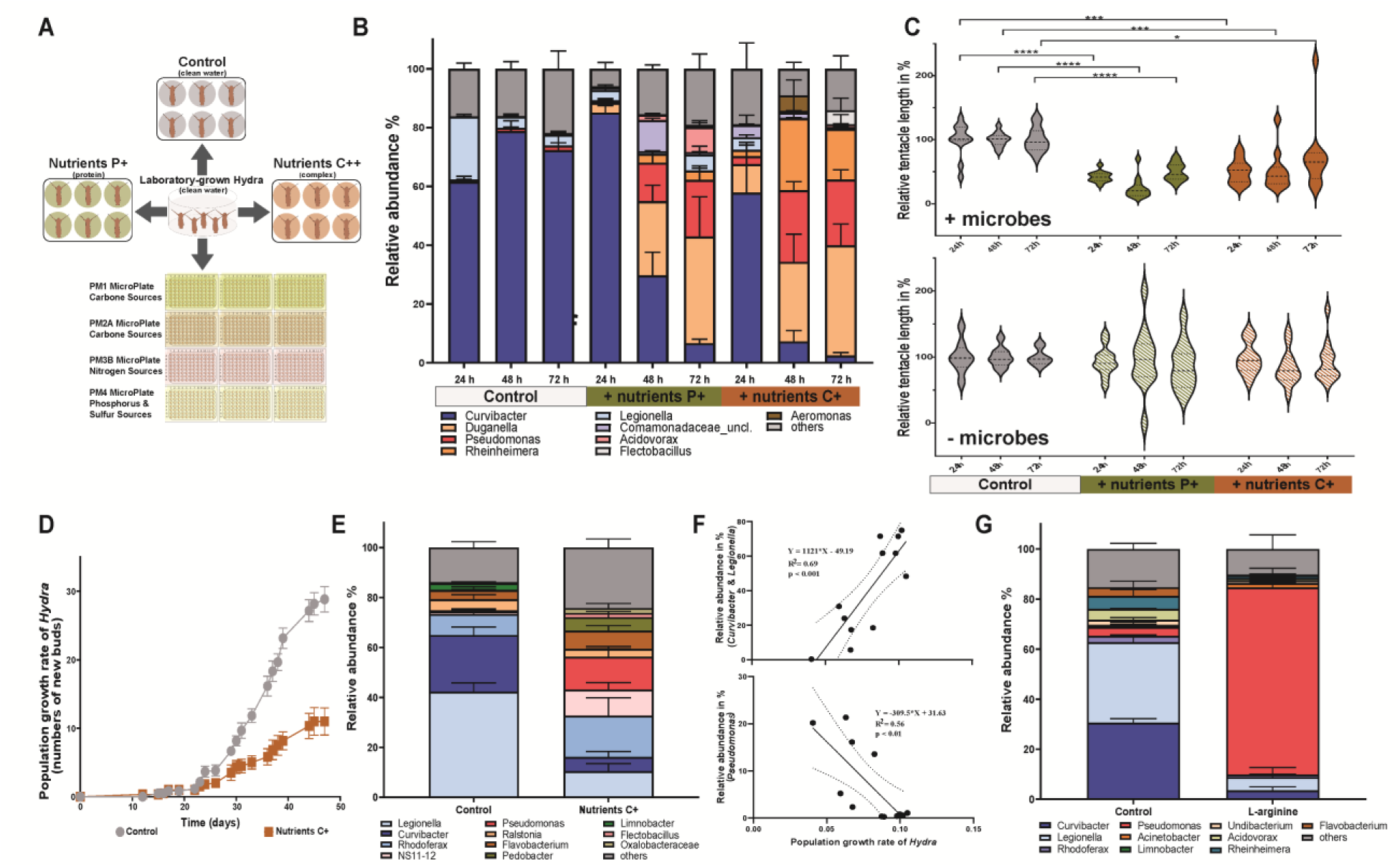
Elevated nutrient levels induce dysbiosis and disease development. (A) Experimental design: Graphic illustration of the experimental setup. Laboratory-grown *Hydra vulgaris* (AEP) were exposed to different nutrient environments, including nutrient-deficient water (control), a protein source (P+) and a complex nutrient source (C+). Individual polyps were also exposed to different compounds in Biolog plates with replication (n=3), allowing analysis of compound-specific effects on microbial community composition. (B) Bar-chart illustrating shifts in microbial community composition based on operational taxonomic units (OTUs) following exposure to to protein source (P+) and complex nutrient source (C+) and nutrient-deficient water. (C) Measurement of polyp tentacle length as an indicator of disease development (+microbes). Dotted lines in violin plots represent the median. One-Way ANOVA with Dunnett’s multiple comparison test compares each group with the control [P > 0.05 = not significant (ns); *P < 0.01; **P < 0.001; ***P < 0.0001; ****P < 0.00001]. Germ-free polyps (-microbes) were used as a control for direct effects of nutrient enrichment in the absence of microbiota. (D) Population growth experiment of *Hydra* exposed to nutrient-deficient water (control) or a nutrient-enriched environment (C+) over a period of 47 days. Population growth was measured by counting bud production per individual poly (n=6). (E) Microbial community composition of the population growth experiment analysed after 47 days of exposure to nutrient-deficient or to nutrient-enriched (C+) environment. (E) *Hydra* population growth rates correlate positively with the relative abundance of *Curvibacter* & *Legionella* (left) and negatively with increasing relative abundance of *Pseudomonas* (right). (F) *Hydra* population growth rates correlate positively with the relative abundance of *Curvibacter* & *Legionella* (top) and negatively with increasing relative abundance of *Pseudomonas* (bottom). (G) Bar-charts illustrating compound-specific shifts in relative abundance of *Hydra* microbiota after 48 h inoculation, highlighting the impact of individual compounds on the microbial community. Data are mean ± s.e.

Most intriguingly, *Hydra* developed a disease phenotype with shortened tentacles(*22*, *23*) under nutrient-enriched environmental conditions. When polyps were exposed to enriched conditions, tentacle length was reduced by 50% within 24 h and remained significantly shorter compared to control polyps until the end of the experiment (Fig. 2, C). This disease phenotype was only observed when polyps were associated with microbes. Tentacle length in germ-free (GF) polyps did not change significantly in nutrient-enriched environments compared to the control (Fig. 2, C).

The protein source (P+) alone induced the same detrimental shifts in microbial community composition, leading to disease development at a lower concentration than the complex medium (C++). In order to ascertain if this was a general phenomenon, we reduced the concentration to 30 mg/l (C+), which corresponds to the dissolved organic load of eutrophic lake environments(*24*) in a secondary experiment. This concentration did not induce visible disease symptoms, but negatively affected the population growth of *Hydra* (Fig. 2, D). Over a period of 47 days, polyps exposed to slightly elevated C+ concentrations produced an average of 12 buds per polyp, in comparison to 32 buds produced by polyps devoid of nutrients. This effect of elevated nutrients on population growth was accompanied by changes in the microbial community (Fig. 2, E). A microbial community analysis conducted at the end of the experiment indicated that the population growth rate of *Hydra* was positively correlated with the abundance of *Curvibacter* and *Legionella*, while negative population growth was associated with increasing *Pseudomonas* abundance (Fig. 2, F).

### Compound-specific fluctuation of the microbial community composition

The results of this study demonstrated two key patterns of the relationship between nutrients and the microbiota: Firstly, the nutrient load and composition of lake water, in particular the carbon-to-nitrogen (C/N) or carbon-to-phosphate (C/P) ratio, significantly influenced the microbial community composition of the host. Secondly, the artificial enrichment of water with a protein or complex nutrient source affected *Hydra*, often to the polyp’s detriment, based on nutrient concentration. These observations strongly suggest that external feeding of the microbiota plays a pivotal role in the development of dysbiosis and disease. To further elucidate the influence of single compounds, rather than complex media, on the composition of the microbiota, we exposed *Hydra* polyps to a variety of different substances in Biolog plates (Fig. 2, A) and characterised the composition of the *Hydra*-associated microbial community after 48 h. Consistent with our previous observations, we detected considerable shifts within the microbial community caused by specific compounds (Fig. S7-10). Interestingly, *Pseudomonas*, which is typically less abundant, with a relative abundance of less than 10 % in nutrient-deficient water, increased to 74 % when *Hydra* was exposed to the amino acid L-arginine (Fig. 2, G). We can conclude that external feeding of the microbial community undermined the competitive advantage of *Curvibacter* in the mucosal environment(*25*). Instead, external feeding favoured microbes that were previously underrepresented in the natural mucus environment. In conclusion, the continuous supply of additional nutrients from the outside dramatically altered the environmental niche occupied by *Hydra*’s microbiota, thereby influencing the composition of its microbial community.

### Harmful infection that originates within the body

#### Changing nutrient-environment conditions turn commensal microbes into pathogens

In all our experiments we observed that disease development and reduced population growth were associated with an increase in the relative abundance of *Pseudomonas*. To gain a deeper understanding of underlying mechanisms, we reduced the complexity of our nutrient source. Based on our observation that the relative abundance of *Pseudomonas* increased in the presence of L-arginine (Fig. 2, H), we mono-colonised polyps with *Pseudomonas alcaligenes* T3 (Fig. S11) isolated from diseased polyps in previous experiments and exposed them to L-arginine (Fig. 3, G). Under these conditions, *Hydra* developed a severe disease phenotype, highlighting the fact that L-arginine metabolism plays a key role in host-microbe interactions (Nüse 2023) and directly affects bacterial virulence and pathogenesis (Gogoi 2016).

**Fig. 3:**
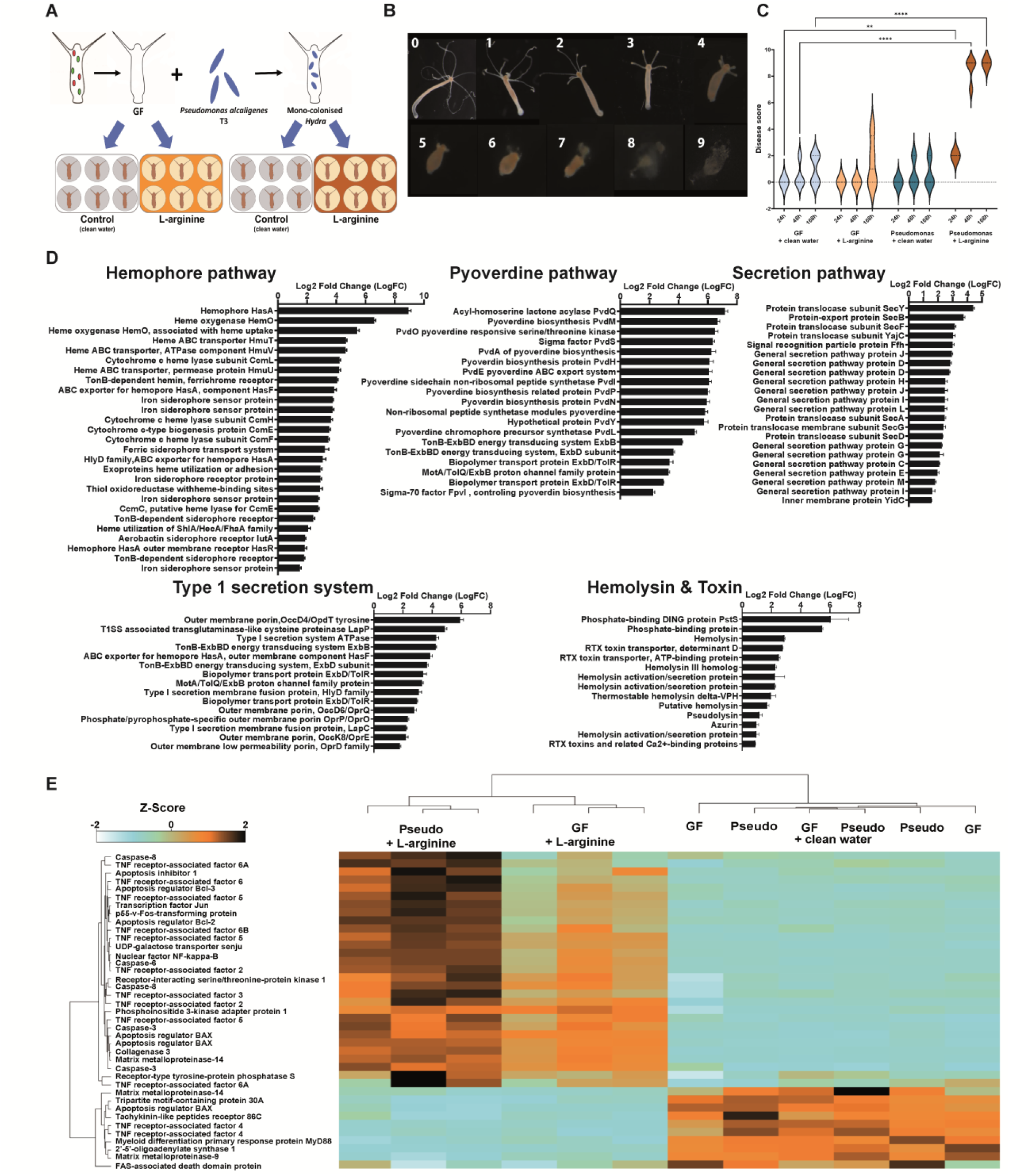
Increased nutrient levels turn a symbiont into a lethal pathobiont. (A) Experimental design: Germ-free (GF) and mono-colonised polyps were exposed to either nutrient-deficient water (control) or L-arginine, with replication (n=6). (B) Disease development in mono-colonised polyps exposed to L-arginine. Disease severity was assessed by grading symptoms. Severity was determined using a numerical scale, starting with a score of 0 for healthy polyps. Reduction in tentacle length and eventual lysis of polyp tissue resulted in scores of 4 to 9. (C) Disease states of polyps were characterised by a scoring system following exposure to either nutrient-deficient water (control) or L-arginine. GF polyps exposed to nutrient-deficient water (light blue), GF polyps exposed to L-arginine (light brown), mono-colonised polyps exposed to nutrient-deficient water (dark blue), and exposed to L-arginine (brown). Dotted lines in violin plots represent the median. One-Way ANOVA with Dunnett’s multiple comparison test compares each group with the control [**P < 0.001; ****P < 0.00001]. (D) Transcriptional changes of *Pseudomonas alcaligenes* T3. Bar-chart illustrating transcriptional changes in *Pseudomonas alcaligenes* T3 when exposed to L-arginine, compared to *Pseudomonas* in nutrient-deficient water. (E) Heatmap of immune response genes: The heatmap specifically focuses on differentially expressed immune genes in *Hydra*, mono-colonised with *Pseudomonas* (Pseudo) and exposed to L-arginine (LArg), belonging to the interleukin and MAPK pathways (based on Z-score values). Germ-free (GF) polyps and nutrient-deficient water (Smed) served as controls. Data are mean ± s.e. (F) Effects of L-arginine supplementation on a *Hydra* symbiont and epithelial cells.

Disease progression was characterized by initial tentacle shrinkage and agglomeration, as well as body length reduction within the first 24 h of exposure. Tentacles then began to disintegrate, and polyps finally dissolved within 48 h. Both mono-colonised polyps exposed to nutrient-deficient water only and GF polyps exposed to nutrient-deficient water or L-arginine did not develop this disease phenotype (Fig. 3, B).

To gain deeper insights into regulatory properties and disease-related transcriptional changes, we sampled *Hydra* for transcriptomic analysis at an early stage of disease development and 24 h post-exposure. RNA sequencing revealed that *P. alcaligenes* T3 colonisation in nutrient-deficient water only had a minor effect on transcription levels in *Hydra* compared to GF polyps (Fig. S12), since only 39 genes were differentially expressed (Fig. S12 and S13).

#### L-arginine induced transition of Pseudomonas from non-pathogenic to pathogenic

Looking at this interaction from a bacterial perspective, L-arginine represents an essential substrate for microbial metabolism and virulence. Supplementation with L-arginine alone resulted in a significant alteration of the transcriptional profile of *P. alcaligenes* T3 (Fig. 3, C), with genes related to iron-acquisition accounting for the majority of differentially regulated transcripts. Pyoverdine biosynthesis and transport systems were upregulated 6-fold when *P. alcaligenes* T3 was exposed to L-arginine instead of nutrient-deficient water. Among other iron scavenging proteins, hemophore HasA was upregulated 9-fold. Type I and Type IV secretion systems were upregulated, while the production of virulence factors such as DING protein, PstS, hemolysins, RTX toxins and azurin was increased (Fig. 3, C). Based on these transcriptional changes, we hypothesize that L-arginine caused a nutrient imbalance through an excessive supply of nitrogen and carbon, forcing *P. alcaligenes* T3 to sequester iron and phosphate to compensate. The increased secretion of iron- and phosphate-binding metabolites, combined with the secretion of various hemolysins and toxins known to disrupt eukaryotic cells, altered the host-bacterial equilibrium, shifting it along the parasite-mutualist continuum(*26*) from a non-pathogenic to a pathogenic state.

#### Transcriptional changes during disease onset in Hydra

This transition towards pathogenicity was already apparent following initial tentacle shrinkage in *Hydra* after 24 h. A transcriptional analysis of polyps mono-colonised with *P. alcaligenes* T3 and exposed to L-arginine, compared to GF polyps in L-arginine, revealed a high proportion of differentially expressed apoptotic, necrotic or immune responsive genes. Among those, several genes belonging to the interleukin and MAPK pathways were upregulated (Fig. 3, D). Upregulation of these pathways likely contributed to the tissue regeneration observed in *P. alcaligenes* T3 induced degradation processes in *Hydra*(*27*).

Another significant group of differentially expressed genes consisted of G-proteins (Fig. S14). Among these, two DKF2 genes, which serve as important regulators of the defence response against gram-negative bacteria(*28*), and two RAS1 genes (activators of MAPK) were upregulated. However, not only genes with an immunostimulatory effect were elevated. Genes that inhibit the release of pro-inflammatory mediators, such as ADRB2, which play an important role in the pathogenesis of asthma(*29*), were upregulated. Additionally, small heat shock proteins such as CRYAB genes, which feature anti-apoptotic activity, were upregulated (Fig. S15). These compensatory responses were likely implemented by *Hydra* to counteract *Pseudomonas*-induced apoptosis and reinforced by the upregulation of three carbonic anhydrases (CAHs). CAHs are involved in the acidification of the human gut environment and are effectors of the innate immune response regulating bacterial infections(*30*)(*31*). These aforementioned innate immune responses and defence mechanisms have an energetic cost. Consequently, polyp shrinkage may be indicative of a reallocation of resources from body mass to host defence, suggesting that this is a potential indicator of disease development.

## Discussion

This study presents evidence for a causal link between environment-induced shifts within the microbiota and the development of disease. Using the model organism *Hydra*, we demonstrated that the host-associated microbiota is variable in its composition. The observed variations in the microbiota are not solely influenced by the host. Instead, it is a complex interplay between the host-epithelial environment, the microbial colonizer pool and conditions of the surrounding environment which select and shape the host-associated microbiota. External feeding of the microbiota annuls the competitive advantage of well-adapted resident microbiota in the mucosal environment(*25*) and abolishes its nutrient blocking capacity, which is important for pathogen protection and colonization resistance(*32*). Instead, other microbes are favored and become dominant by external feeding, despite being underrepresented in the natural mucus environment. Overfeeding of the microbiota, as postulated by the “overfeeding hypothesis”(*11*), allows for uncontrolled microbial growth and can transform neutrally associated microbes into pathogens, depending on the nutrient composition. Humans are associated with a wide range of commensal or beneficial microbes. However, whether these microbes are genuinely harmless depends on environmental conditions, which shift microbes along the parasite-mutualist continuum(*26*).

## Acknowledgments

We appreciate the support in the field experiment by the Landesamt für Landwirtschaft, Umwelt und ländliche Räume (LLUR), in particular Mrs. Elisabeth Wesseler and Mrs. Angelika König. The Landeslabor Schleswig-Holstein, in particularly Mrs. Gerda Rügner, for carrying out the lake water nutrient analysis. We acknowledge the sequencing support by the Kiel Institute for Clinical Molecular Biology (IKMB) and Prof. Dr. Dr. h.c. Thomas C.G. Bosch for giving us the opportunity to conduct this work in his laboratory. For utilising their Galaxy EU server, we thank the University of Freiburg (Germany), funded by the Collaborative Research Centre 992 Medical Epigenetics (DFG grant SFB 992/1 2012) and the German Federal Ministry of Education and Research BMBF grant 031 A538A de.NBI-RBC. For her feedback on a previous version of the manuscript we thank Katrin Hammerschmidt.

## Author Contributions

T.L. conceptualization, formal analysis, investigation, data curation, writing – original draft preparation, review and editing, visualization, supervision, project administration, funding acquisition. L.U. and F.M.W. investigation (mono-colonisation experiments). L.U. formal analysis (RNA-seq data), visualization, writing – review and editing.

M.W. investigation (Biolog and mono-colonisation experiments), formal analysis. T.H. investigation (nutrient exposure experiments), formal analysis. L.X.S. investigation (genome sequencing), formal analysis (genome annotation). E.V.H. investigation (relocation, population growth, nutrient exposure, Biolog experiments). P.D. conceptualization, investigation, writing – review and editing.

## Data availability

All data supporting the findings of this study are available within the paper and its Supplementary Information. Sequence data is deposited in National Center for Biotechnology Information and is available under the project ID PRJNA997121.

## Supplementary Materials

### Materials and Methods

#### Long-term laboratory *Hydra* culture conditions

*Hydra vulgaris* (AEP)(*33*) was cultured under constant laboratory conditions in nutrient-deficient water (s-medium: 0.28 mM CaCl_2_, 0.33 mM MgSO_4_, 0.5 mM NaHCO_3_ and 0.08 mM KCO_3_) at 18 °C water temperature and a 12 h light-dark cycle, according to the standard procedure(*34*).

#### I: Exposing laboratory-grown *Hydra* to natural lake environments

##### Field experiment

Laboratory-grown *Hydra vulgaris* (AEP) polyps with their associated bacterial community were exposed to different lake environments for a period of up to 7 days. *Hydra* polyps were relocated to three mesotrophic lakes (Schluensee, Selenter See and Pluf3see) and three eutrophic lakes (Lanker See, Postsee and Tresdorfer See) (Fig. S1). For polyp retrieval, 15 polyps per replicate were transferred into 50 ml falcon tubes (n=5). The openings were sealed with gauze (150 µm) to allow exchange with the surrounding environment and influx of bacteria, while preventing immigration of zooplankton and polyp loss. Tubes containing polyps were tied together and deployed at 30 cm water depth. After 1, 4 and 7 days one polyp per treatment and replicate (n=5) was collected. Polyps were transferred separately to 1.5 ml tubes, washed twice in 500 µl sterile s-medium and stored at -20 °C until DNA extraction.

##### Laboratory experiment

In parallel to the field experiment, a similar experiment was carried out under controlled laboratory conditions at 18 °C water temperature and a 12 h light-dark cycle. Lake water was collected from different lake environments (see above) and filtered through 1.5 µm to remove larger organisms and particles. *Hydra* polyps were transferred separately into 6-well plates. 5 polyps per replicate (n=6) were exposed to the 6 different lake water environments. On days 1, 4 and 7, one polyp per replicate was collected in a 1.5 ml tube, washed twice in 500 µl sterile s-medium, and stored at - 20 °C until DNA extraction.

##### Exposure of laboratory-grown Hydra to lake nutrients

Lake water was filtered at 0.02 µm in order to remove all bacteria and phages to test the effect of the chemical environment on *Hydra’s* microbial community composition. The nutrient composition of the lake water, including dissolved organic carbon (DOC), total nitrogen and total phosphate, was analysed by the Landeslabor Schleswig-Holstein and LLUR (Table S1). Water was changed daily. After 7 days of exposure, polyps were collected and prepared for DNA extraction (see below).

#### II: Nutrient manipulation of the environment

##### Exposure to complex nutrients

The effect of elevated nutrients on *Hydra* host-microbe homeostasis was tested by adding either a protein source (Envir1) or a complex nutrient source (Envir2) to nutrient-deficient water (s-medium). Envir1 one was prepared by adding 0.05 mg/ml peptone, while Envir2 was made using bacterial culture medium R2A (ROTH) at a final concentration of 0.3 mg/ml. To control for the effect of nutrients on the eukaryotic host in the absence of associated bacteria, germ-free (GF) and wildtype (wt) polyps were exposed separately to nutrient-enriched water conditions. Polyps exposed to nutrient-deficient water (s-medium) served as controls. GF *Hydra* were generated by exposing animals to an antibiotic cocktail containing 50 µg/ml of ampicillin, rifampicin, streptomycin, spectinomycin and neomycin for a period of two weeks(*23*). Antibiotic solutions were exchanged every second day. To remove remaining antibiotics, animals were transferred to antibiotic-free, sterile s-medium for three days after antibiotic treatment. Sterility was confirmed by negative bacterial 16S PCR(*13*). The effect of nutrients on the host-associated microbial community was analysed every 24 h (n=5). Polyps were washed three times and finally homogenised in 500 µl sterile s-medium. 100 µl were plated onto R2A agar (ROTH) plates to establish a bacterial culture collection that served as a reservoir of potential bacterial pathogens. 200 µl were used for DNA extraction using the DNA Blood and Tissue Kit (Qiagen). Microbial community composition was determined by rRNA gene amplicon sequencing (see below).

##### Evaluation of disease symptoms

Disease development in *Hydra* is typically measured by scoring morphological changes (*22*, *23*). As disease development starts with shrinking tentacles, we measured tentacle length every 24 h for a period of 4 days (n=12) to access early signs of morphological changes. Polyps were photographed using a binocular microscope and length estimates were made using ImageJ 1.50i (*35*). Severe disease states were assessed through symptom grading. The severity was determined using a numerical scale, starting with a score of 0 for healthy polyps, progressing to a reduction in tentacle length and eventual lysis of polyp tissue (scales 3 to 1). Total tissue degradation, accompanied by the loss of polyp body shape, was scored as 9.

##### Population growth experiment

To test the effect of elevated nutrient concentrations corresponding to eutrophic natural lake environments, *Hydra* polyps were individually transferred to 6-well plates (n=6) containing 0.03 mg/ml R2A. Population growth was estimated by counting bud production over a period of 47 days. At the end of the experiment, polyps were removed, washed in sterile s-medium and stored at -20 °C until DNA extraction.

##### Biolog experiment

In total, l,l52 *Hydra vulgaris* (AEP) polyps were washed and kept in sterile s-medium for 3 days without feeding prior to exposure to Biolog plates. The compounds in the Biolog plates (PMl MicroPlate™ Carbon Sources, PM2A MicroPlate™ Carbon Sources, PM3B MicroPlate™ Nitrogen Sources and PM4A MicroPlate™ Phosphorus and Sulphur Sources with replication (n=3)) were dissolved in sterile s-medium and diluted 20-fold. Polyps were randomly distributed into Biolog plates, placing one polyp per well. After 48 h of exposure, polyps were removed, washed twice in sterile s-medium, and then stored at -20 °C until DNA extraction.

##### Recolonisation experiment

Germ-free *Hydra* were mono-colonised with *Pseudomonas alcaligenes* T3, so that 5,000 CFUs/ml were added into the surrounding water of GF polyps and incubated for one day. Non-attached bacteria in surrounding medium were removed by washing polyps in sterile s-medium. Mono-colonised polyps and GF polyps (n=5) were exposed to either L-arginine or s-medium. Disease development was evaluated by scoring morphological changes(*22*, *23*).

##### Microbial community analysis

200 µl of homogenised polyps (see above) were used for DNA extraction (DNA Blood and Tissue Kit (Qiagen). Microbial community composition was determined by amplicon sequencing of the variable region V1-V2 of the 16S rRNA gene. We used the following primers: Forward primer 27F: (5′-AATGATACGGCGACCACCGAGATCTACAC XXXXXXXX TATGGTAATTGT AGAGTTTGATCCTGGCTCAG-3′). Reverse primer 338R: (5′-CAAGCAGAAGACGGCATACGAGAT XXXXXXXX AGTCAGTCAGCC TGCTGCCTCCCGTAGGAGT-3′). Primers contained the Illumina adapter p5 (forward) and p7 (reverse) and unique MIDs (designated as XXXXXXXX) to label each PCR product. PCR reactions were performed in duplicate using Phusion Hot Start DNA Polymerase (Finnzymes, Esppoo, Finnland). PCR cycling conditions were: 98 °C for 30 s, 30 x [98 °C – 9 s, 55 °C – 30 s, 72 °C – 90 s], 72 °C – 10 min. PCR products were combined and purified by using the MinELute Gel Extraction Kit (Qiagen) after agarose gel electrophoresis. Sequencing was performed on the Illumina MiSeq platform at the sequencing facility of the Kiel Institute for Clinical Molecular Biology (IKMB). Sequencing data was analysed using the MOTHUR packages(*36*) according to the MiSeq SOP(*37*). In summary, MiSeq paired-end reads were assembled and quality controlled resulting in 12,028 sequences per sample. Sequences were grouped into operational taxonomic units (OTUs) using a 97 % similarity threshold. Sequences were aligned to the SILVA 128 Database and taxonomically classified by the RDP classifier. Multidimensional scaling analysis of OTU abundance data based on Bray-Curtis similarity was performed by the Primer software Version 7.0.l3 (Primer-E)(*38*). Similarities between different treatment groups were analysed by ANOSIM (analysis of similarity). ANOSIM pairwise test calculated R-values close to 1 indicate that group similarity is higher within than in between different groups. Raw data are deposited in the Sequence Read Archive (SRA) and are available under project ID PRJNA997l2l-SUBl3698683 (lake environment) - SUB13699056 (complex nutrients) - SUBl3704l20 (Biolog) - SUBl38l6039 (Biolog) - SUB13698848 (population growth).

##### Pseudomonas alcaligenes 13 genome sequencing and annotation

The Nextera XT kit (Illumina) was used for library preparation. Bacterial DNA was 2 x 150 bp paired-end sequenced on a NovaSeq platform (Illumina) at the IKMB in Kiel. Raw Illumina reads were adapter-, quality-trimmed and filtered using BBTools v38.96(*39*) and fastp v0.23.2(*40*).

A long-read library was made using the Rapid Sequencing Kit (SQK-RAD004) and sequenced on the MinION (Oxford Nanopore Technologies, Oxford, UK) with a Flongle flow cell (FLO-FLG00l). The super-accurate model of Guppy (Oxford Nanopore Technologies plc. Version 6.2.l+6588ll0, dna_r9.4.l_450bps_sup) was used for base-calling and reads were adapter-trimmed with Porechop v0.2.4(*41*). A hybrid genome assembly, using short and long-reads, was generated with Unicycler v0.5.0.(*42*), and its completeness evaluated with CheckM v1.2.2(*43*) and BUSCO v5.4.3(*44*). The genome sequences of *Pseudomonas alcaligenes* T3 is available under project ID PRJNA997l2l-SUBl3807467.

##### Phylogeny

Taxonomic classifications were performed with GTDB-Tk v.2.2.6(*45*) using the classify and denovo workflow with *Pseudomonas* reference genomes from the Genome Database Taxonomy (GTDB, r207)(*45*). GTDB-Tk identified T3 as *P. alcaligenes* (GCF_000474255.1) based on average nucleotide identity (ANI) of 98.64 %. A denovo pipeline was used to calculate a phylogenetic tree and to confirm the genome placement and identification. For this, genomes from *Pseudomonas* group F and O (as outgroup) were used in the alignment.

##### Hydra RNA extraction

GF and mono-colonised polyps with *Pseudomonas alcaligenes* T3 were exposed to either s-medium or L-arginine (n=3). In total, 15 polyps per replicate (n=3) were incubated under these conditions for 24 h. Polyps were collected and dissolved in 750 µL trizol at room temperature and stored at -80 °C. Upon thawing, 250 µL of chloroform was added, samples were mixed and incubated at room temperature for 5 min. Samples were centrifuged at l2,000 g for l5 min at 4 °C. The upper phase was transferred into a new tube, mixed with 1x volume of cold ethanol and transferred into Spin Cartridges. Subsequent purification steps were conducted according to the PureLink™ RNA Mini Kit (ThermoFisher Scientific), with the exception that we doubled all washing steps. RNA was eluted into 35 µL RNase-free water and stored at -80 °C prior to sequencing.

##### Hydra RNA-seq and data analysis

Strand-specific cDNA libraries were prepared using TruSeq adapters and sequenced (paired-end) via NovaSeq 6000 S4 PE150 XP RNA. Sequences were analysed according to Batut *et al.*(*46*). In brief, we used Cutadapt(*47*), Trimmomatic(*48*), FastQC (Andrews, n.d.) and MultiQC(*49*) for quality control. We used RNA Star(*50*) to map reads to the *Hydra vulgaris* (AEP) genome provided by Cazet *et al.*(*51*). Featurecounts(*52*) was used to count the reads. Finally, we conducted differential gene expression analysis via Deseq2(*53*). The raw data is deposited at the Sequence Read Archive (SRA) and is available under project ID PRJNA997l2l.

##### Bacterial RNA-seq

*Pseudomonas alcaligenes* T3 was exposed to either s-medium or L-arginine with replication (n=4). After 12 h of exposure, bacterial cells were harvested by centrifugation and bacterial RNA extracted according to the method described above. Illumina stranded total RNA preparation was used for ribosomal removal and cDNA library preparation. Sequencing was conducted on a NovaSeq 6000 platform (Illumina). Sequence reads were trimmed and adapters removed using Trimmomatic-0.36(*48*). Trimmed and quality controlled reads were mapped separately against the genome of *P. alcaligenes* T3 using the computer software Bowtie2(*54*) and SAM tools(*55*). Coverage was calculated and normalized to the housekeeping gene rpoS(*56*). Raw data is deposited in the Sequence Read Archive (SRA) and is available under project ID PRJNA997l2l-SUBl38249l3.

###### Data availability

All data supporting the findings of this study are available within the paper and its Supplementary Information. Sequence data is deposited in National Center for Biotechnology Information and is available under the project ID PRJNA997l2l.

### Supplementary Figures

**Fig. S1.**
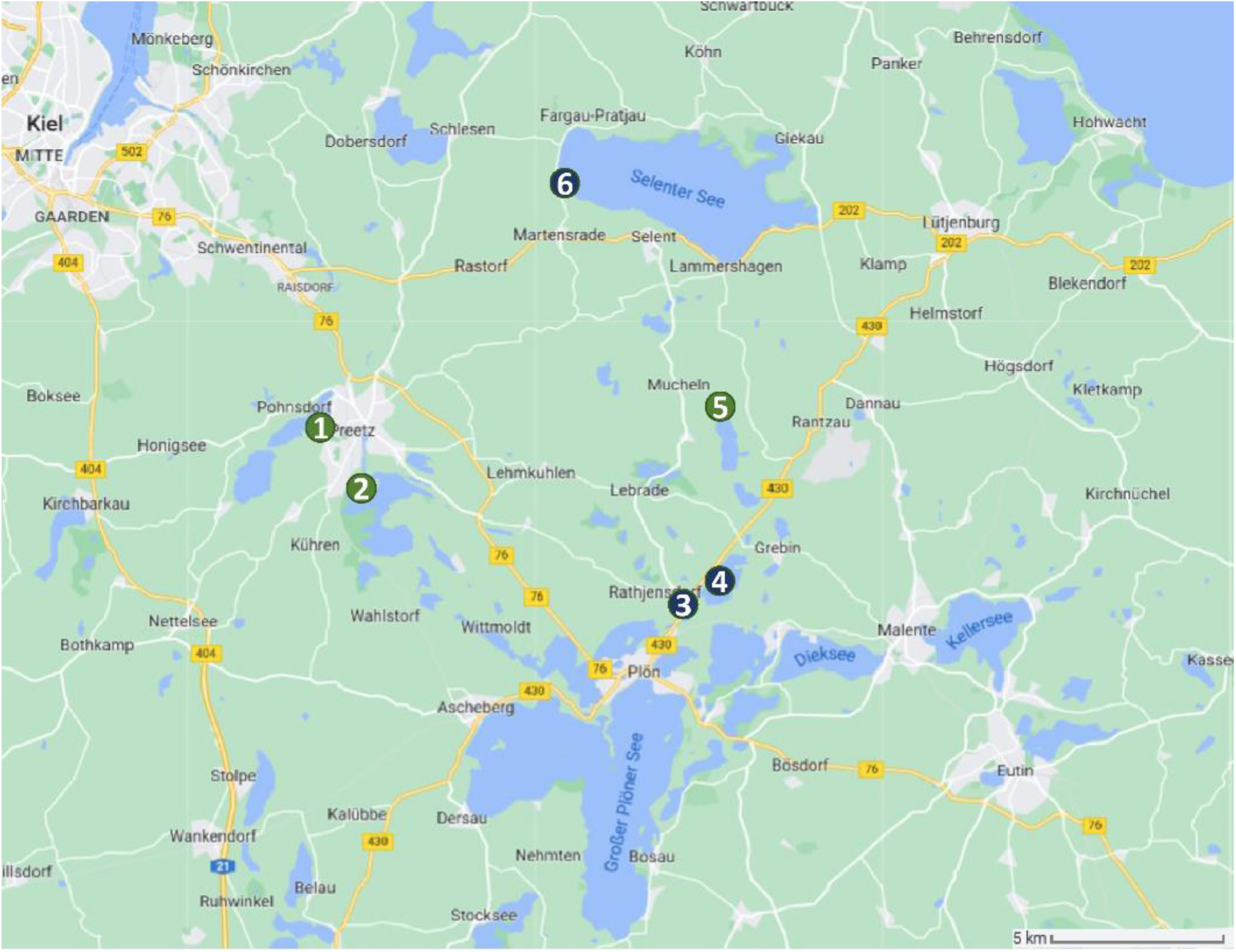
Experimental sites. (1) Postsee=**PO** (eutroph), 242ll Pohnsdorf 54°14’05.5"N l0°l5’46.3"E; 54.234867, 10.262865; (2) Lanker See=**LA** (eutroph), 242ll 54°l3’l3.3"N l0°l7’53.l"E; 54.220360, 10.298077; (3) Pluf3see=**PL** (mesotroph), 24306 Rathjensdorf 54°ll’02.4"N 10°26’32.9"E; 54.l84006, l0.442480; (4) Schluensee=**SC** (mesotroph), 54°ll’08.2"N 10°27’l6.l"E; 54.l85604, l0.454465; (5) Tresdorfer See=**TR** (eutroph), 54°l3’5l.5"N l0°27’55.7"E; 54.230980, 10.465476; (6) Selenter See=**SE** (mesotroph), 54°17’54.0"N 10°25’08.1"E; 54.298335, l0.4l89l5.

**Fig. S2.**
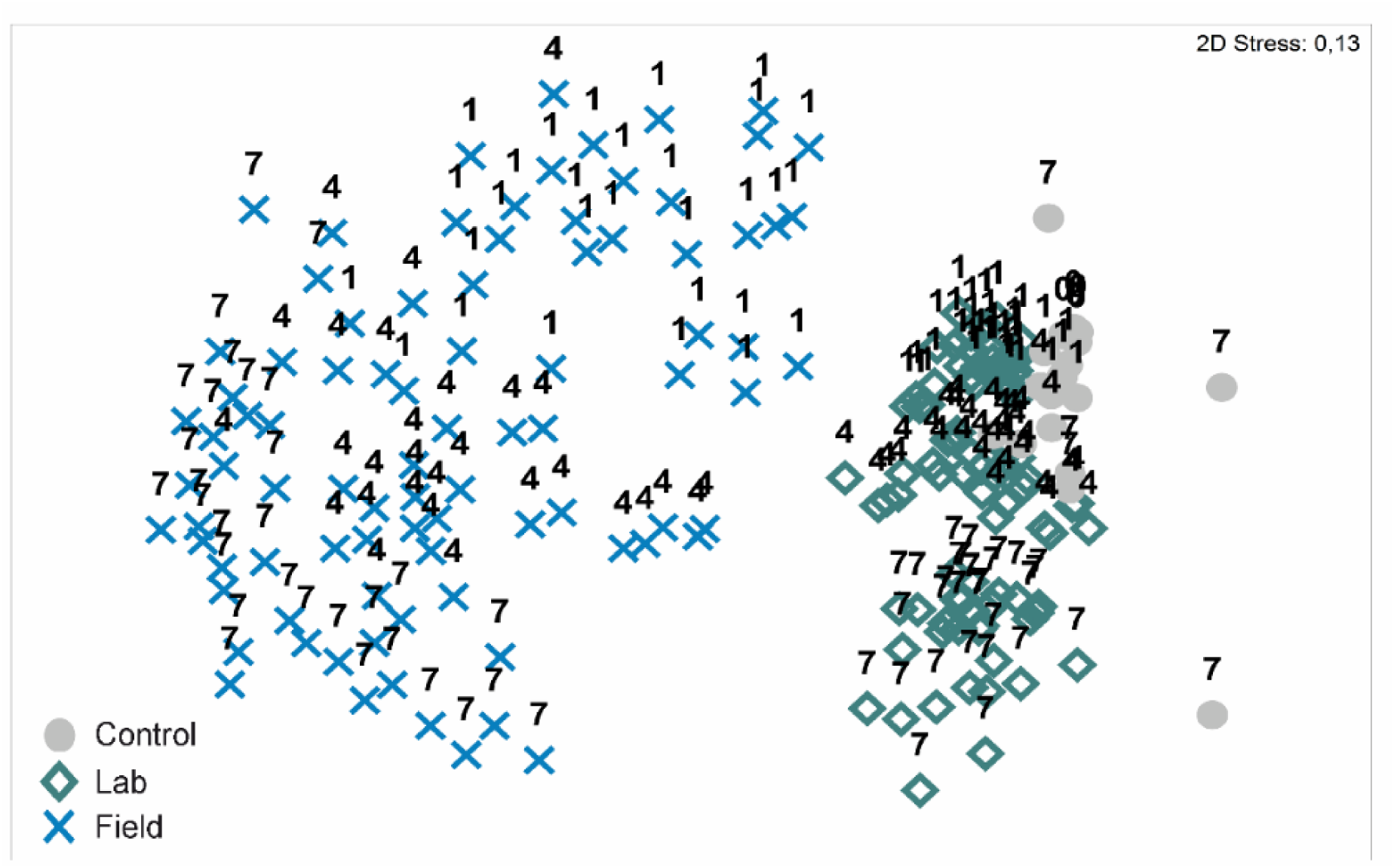
nMDS analysis of *Hydra*-associated microbial community composition exposed to different lake water environments (laboratory and field) based on 16S rRNA gene amplicon sequencing. Nutrient-deficient water (s-medium) was used as control in the laboratory. Microbial community composition of control polyps was significantly differed from polyps exposed to different lake water environments (Permanova P<0.00l).

**Fig. S3.**
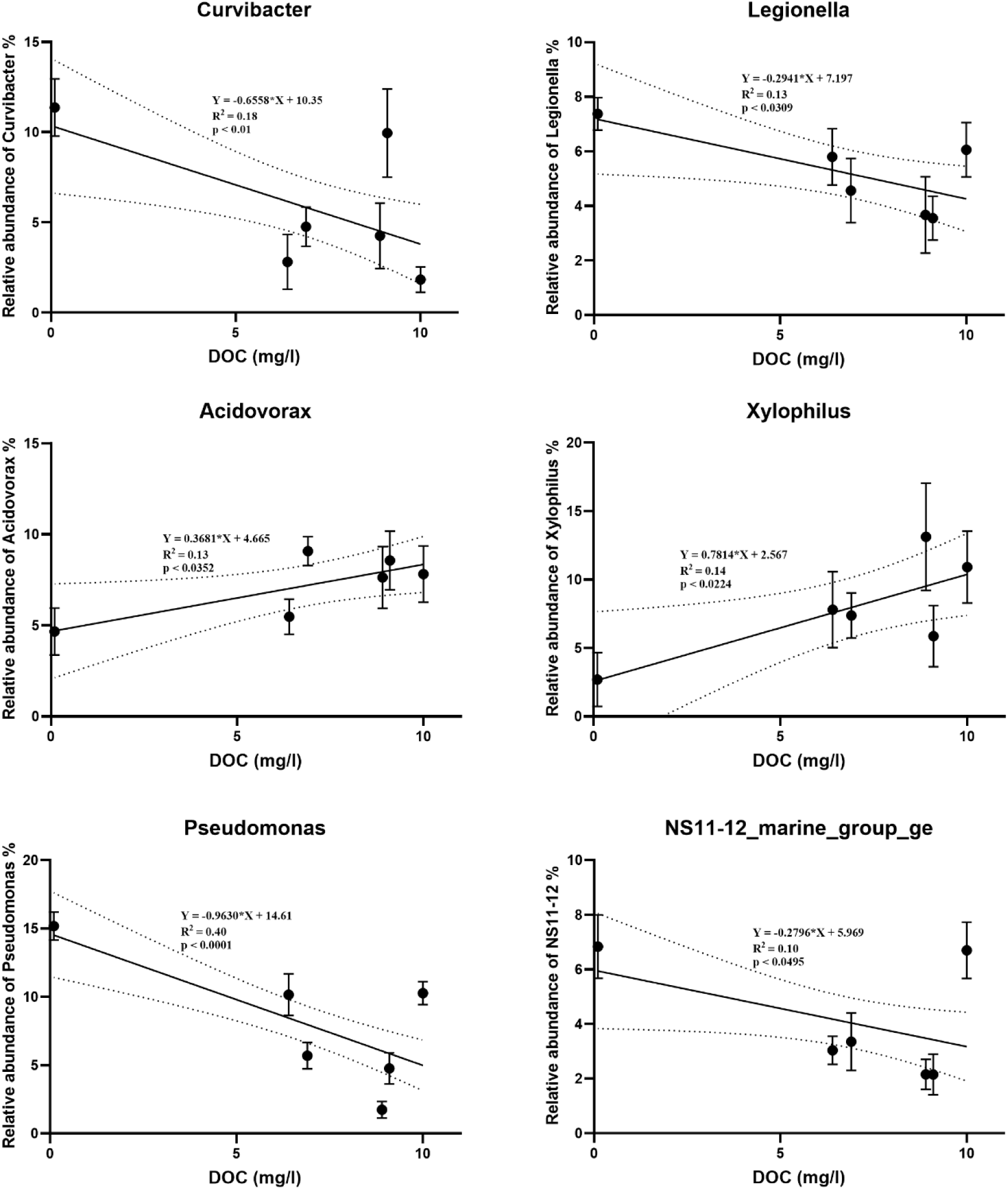
Simple linear regression between the relative abundance of bacteria and dissolved organic carbon (DOC).

**Fig. S4.**
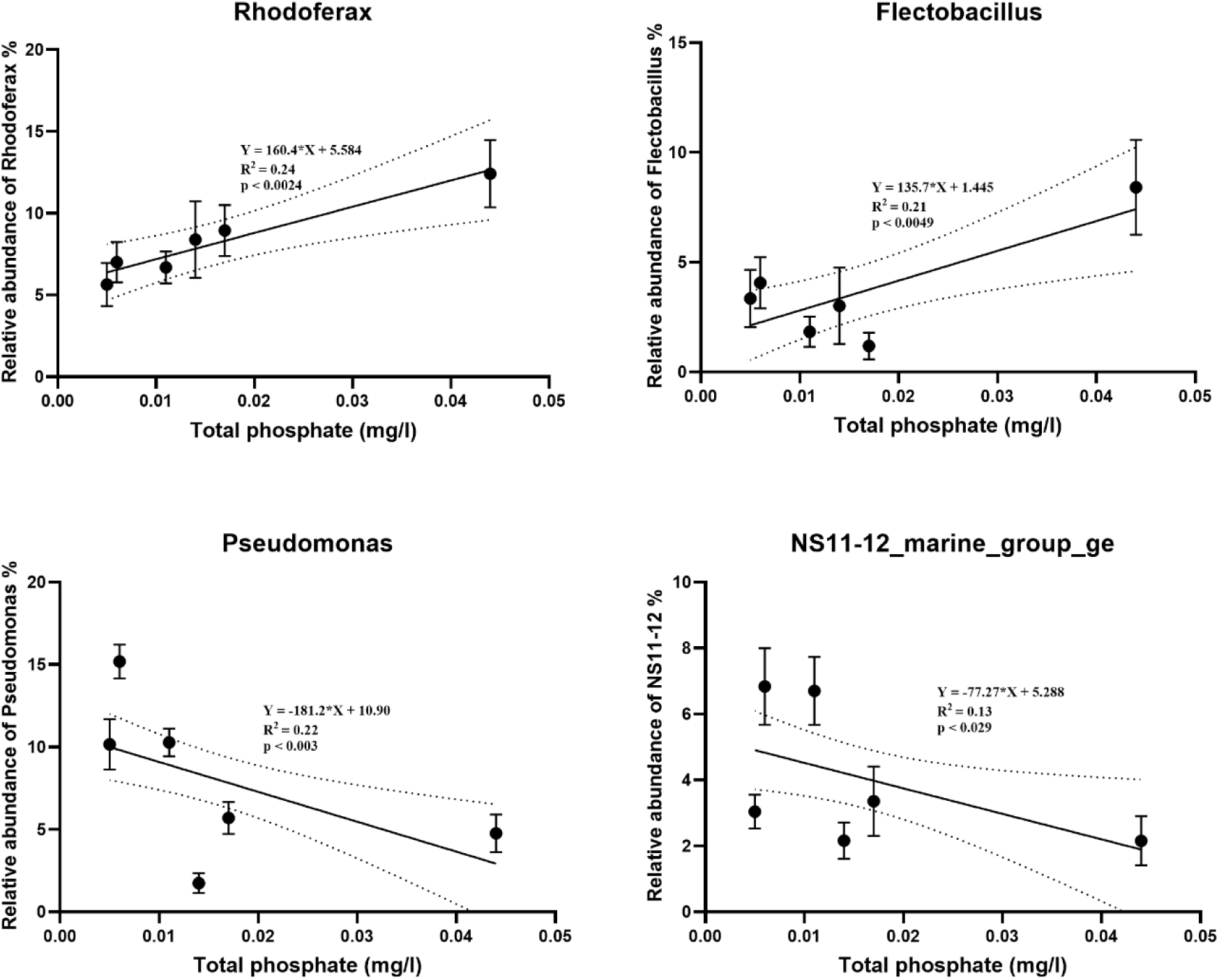
Simple linear regression between the relative abundance of bacteria and total phosphate.

**Fig. S5.**
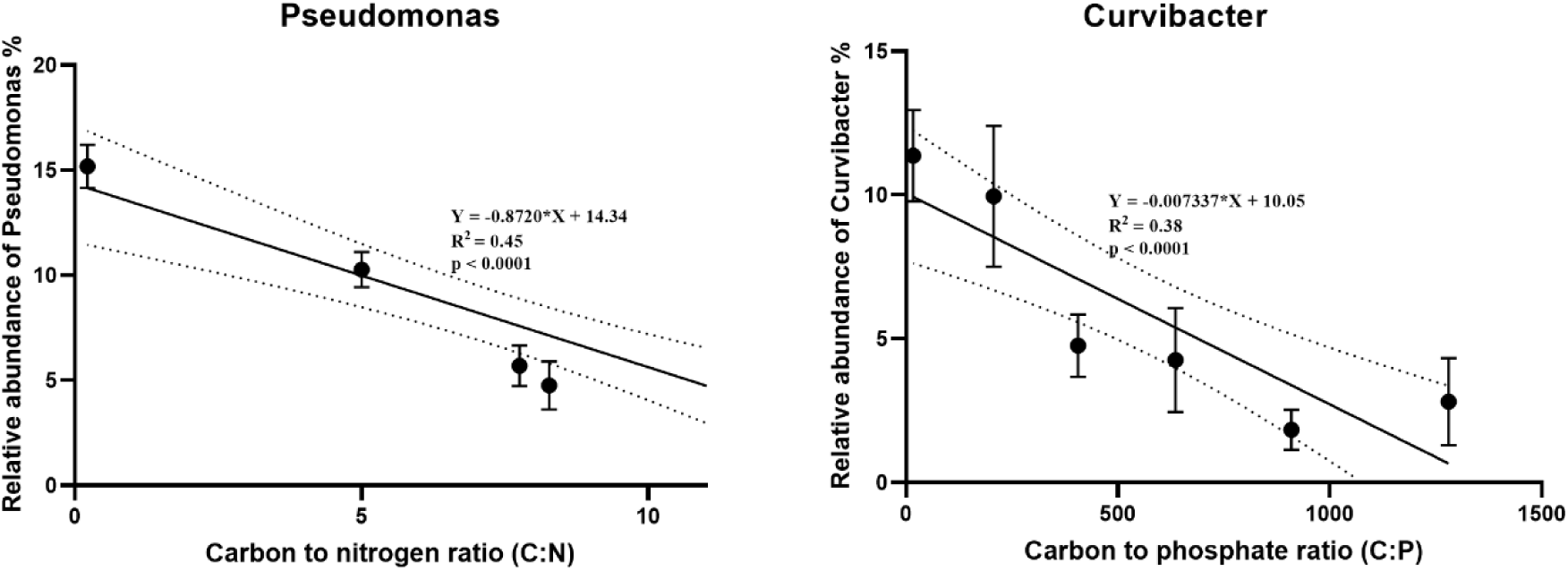
Simple linear regression between the relative abundance of bacteria and nutrient ratios (C/N and C/P).

**Fig. S6.**
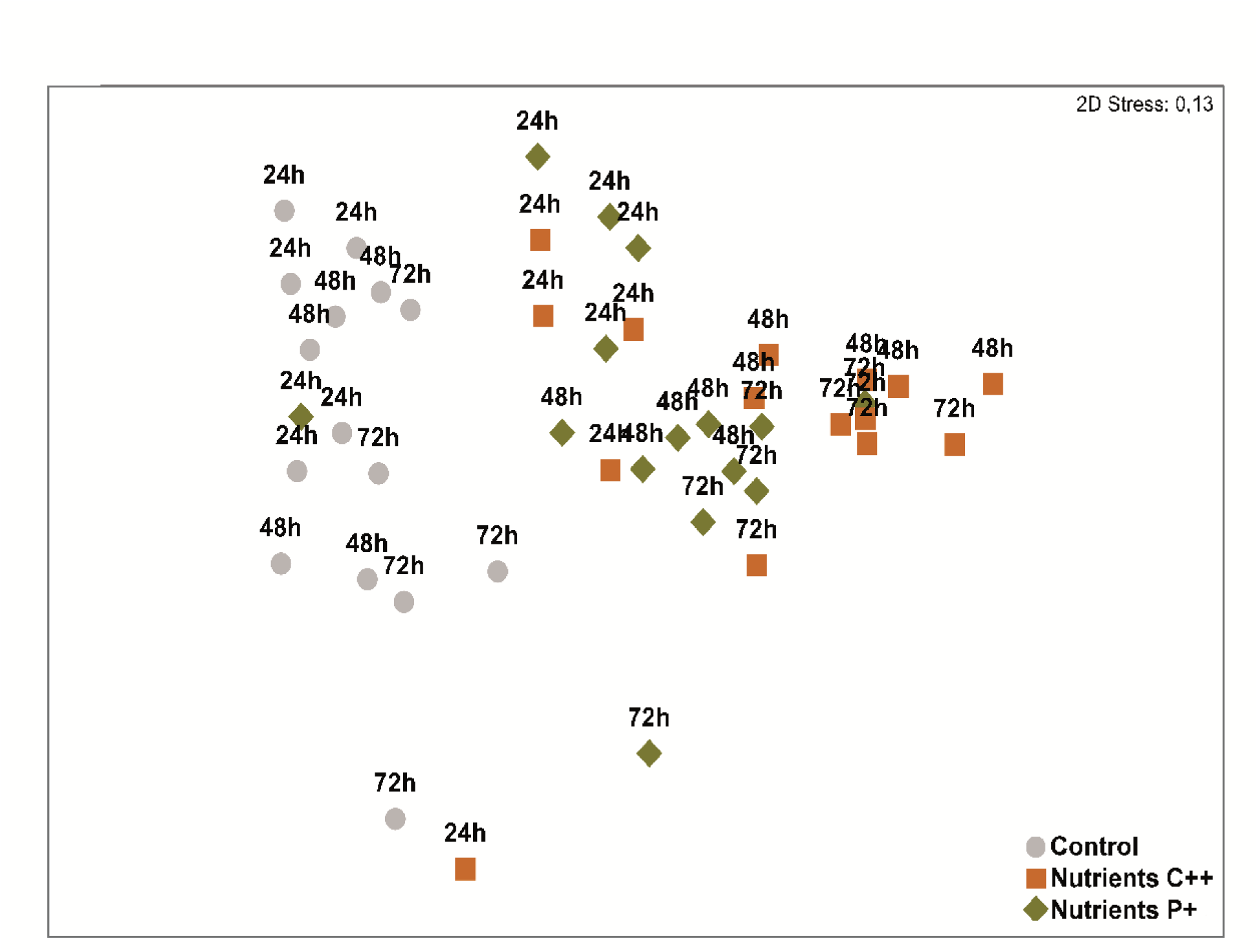
External feeding of mucus-associated bacteria on *Hydra* altered the microbial community composition. Nonmetric multidimensional scaling (nMDS) analysis of 16S rRNA gene amplicon sequencing data revealed two distinct clusters. Polyps exposed to nutrient-enriched environments clustered separately from control polyps in nutrient-deficient water (grey circles). The microbial community composition of control polyps was significantly differed from polyps exposed to nutrient C++ (ANOSIM Pairwise-Test: R Statistic=0.776, P=0.001) and nutrient P+ (ANOSIM Pairwise-Test: R Statistic=0.832, P=0.00l).

**Fig. S7.**
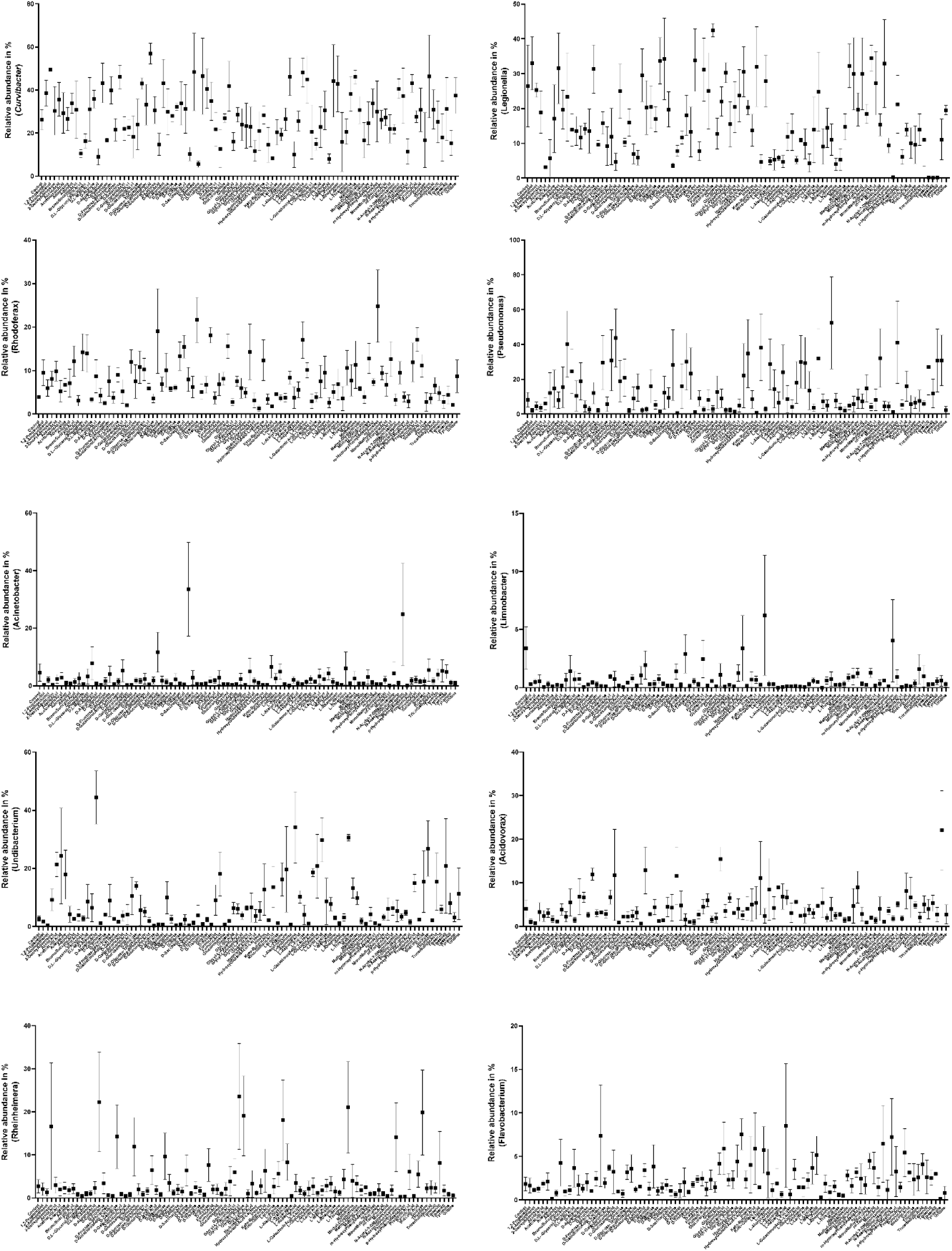
Bacterial community shifts of *Hydra*-associated bacteria following a 48 h exposure to different compounds of the PM1 MicroPlate™ (Carbon Sources). Data are mean ± s.e.

**Fig. S8.**
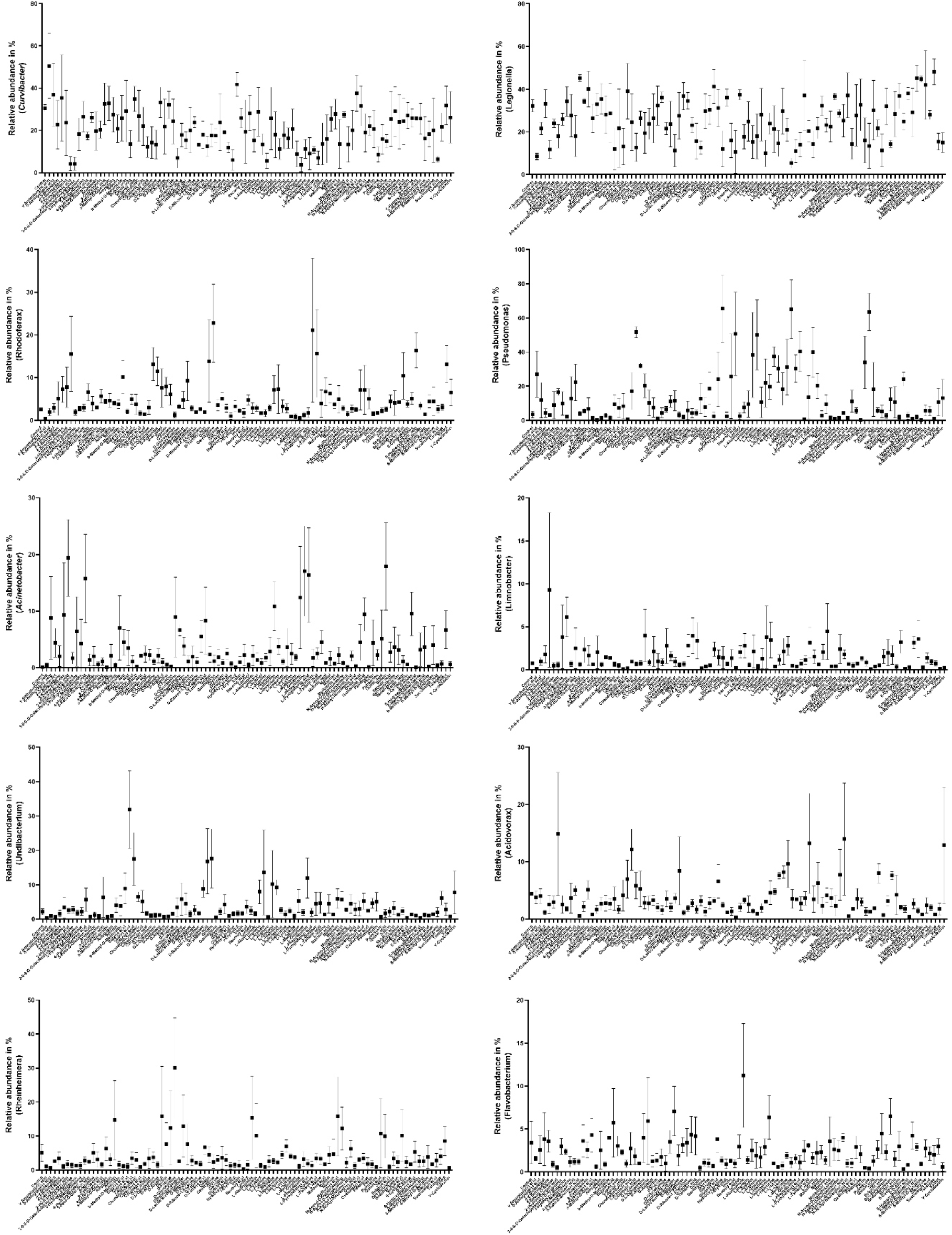
Bacterial community shifts of *Hydra*-associated bacteria following a 48 h exposure to different compounds of the PM2A MicroPlate™ (Carbon Sources). Data are mean ± s.e.

**Fig. S9.**
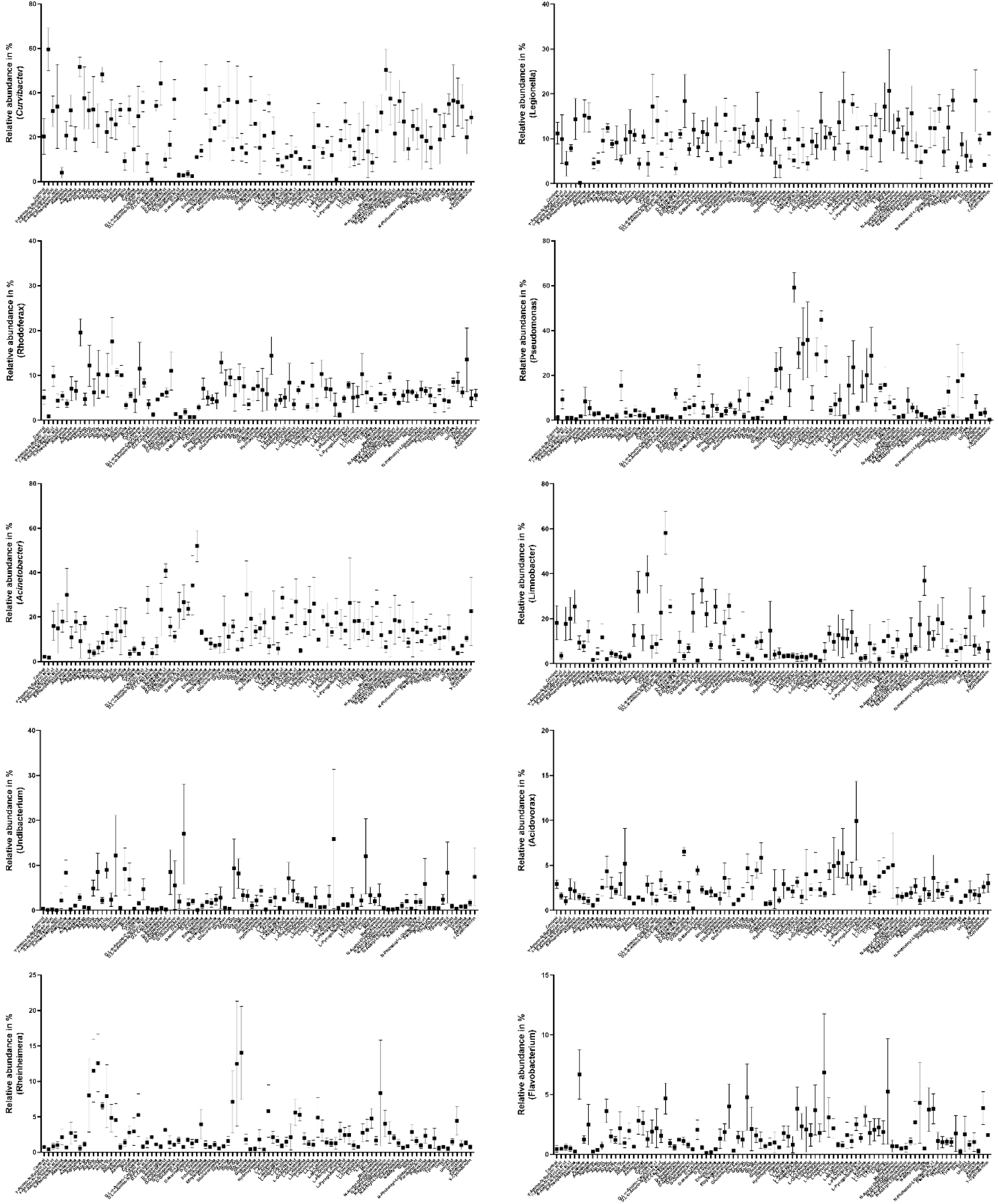
Bacterial community shifts of *Hydra*-associated bacteria following a 48 h exposure to different compounds of the PM3 MicroPlate™ (Nitrogen Sources). Data are mean ± s.e.

**Fig. S10.**
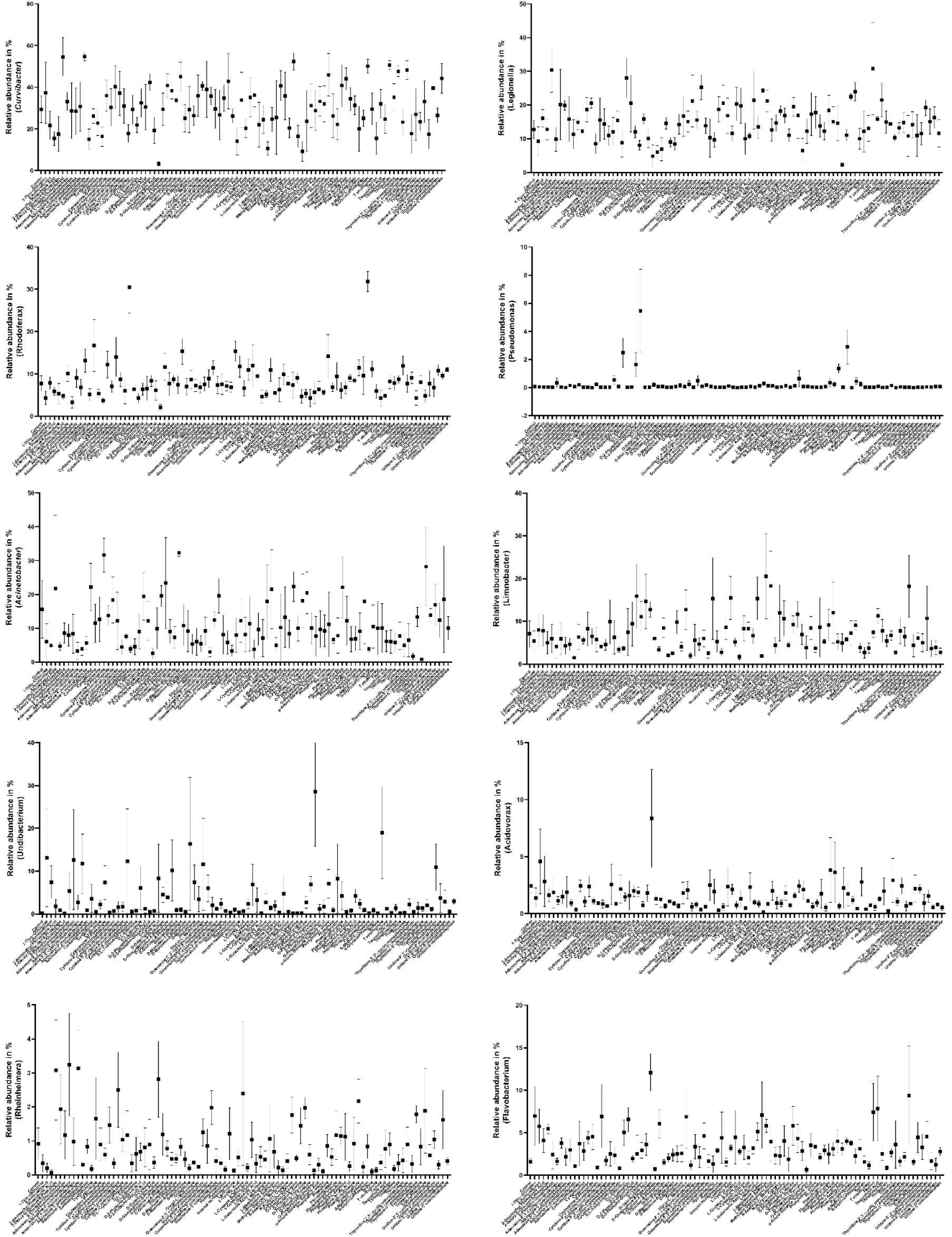
Bacterial community shifts of *Hydra*-associated bacteria following a 48 h exposure to different compounds of the PM4A MicroPlate™ (Phosphorus and Sulphur Sources). Data are mean ± s.e.

**Fig. S11.**
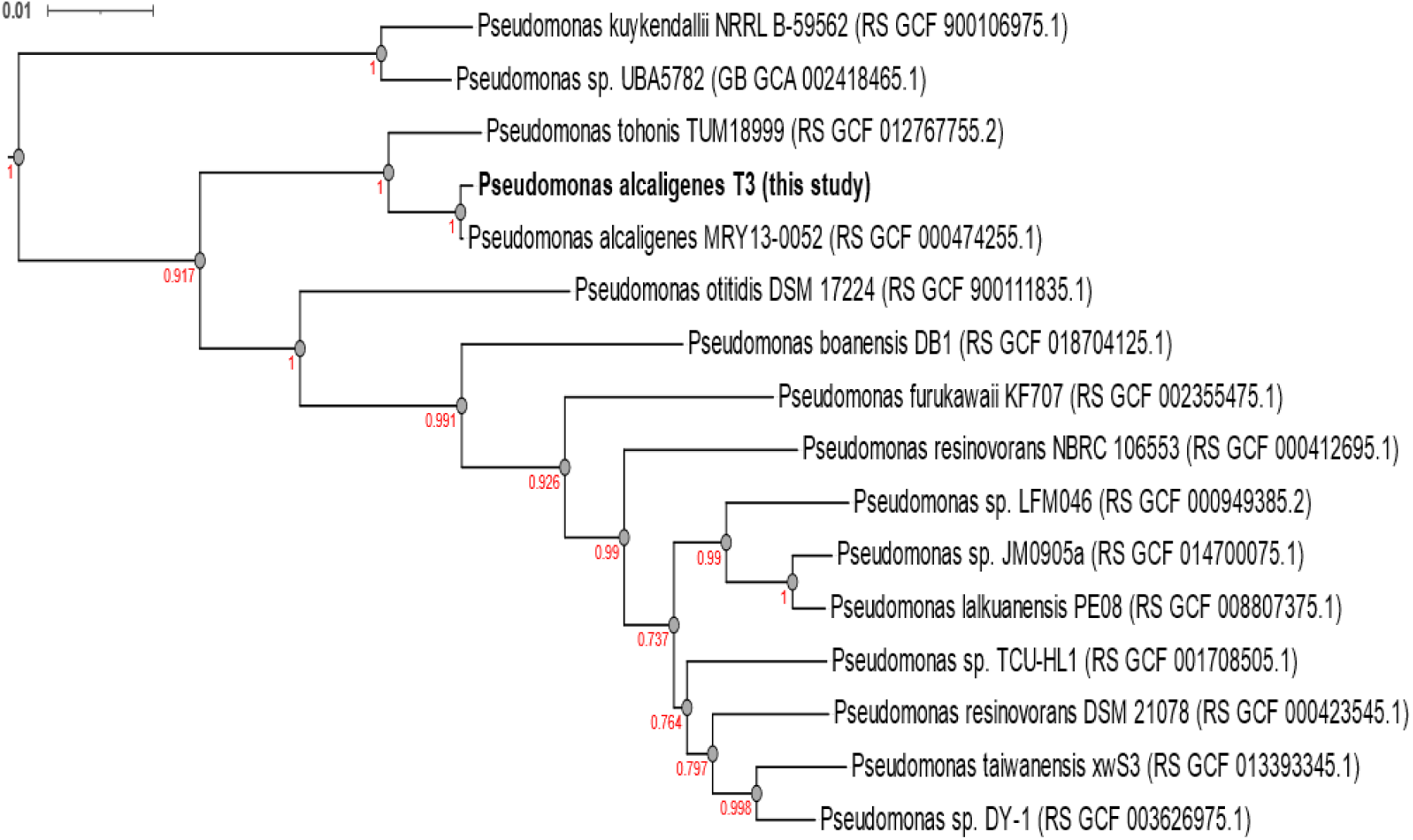
Phylogenetic tree: Genome phylogeny of *Pseudomonas alcaligenes* T3 with reference genomes of *Pseudomonas* group F (and group O used as outgroups) in the Genome Taxonomy Database (GTDB). A maximum-likelihood phylogenetic tree is constructed based on 120 bacterial single-copy marker proteins. The bar indicates XXX substitutions per nucleotide position.

**Fig. S12.**
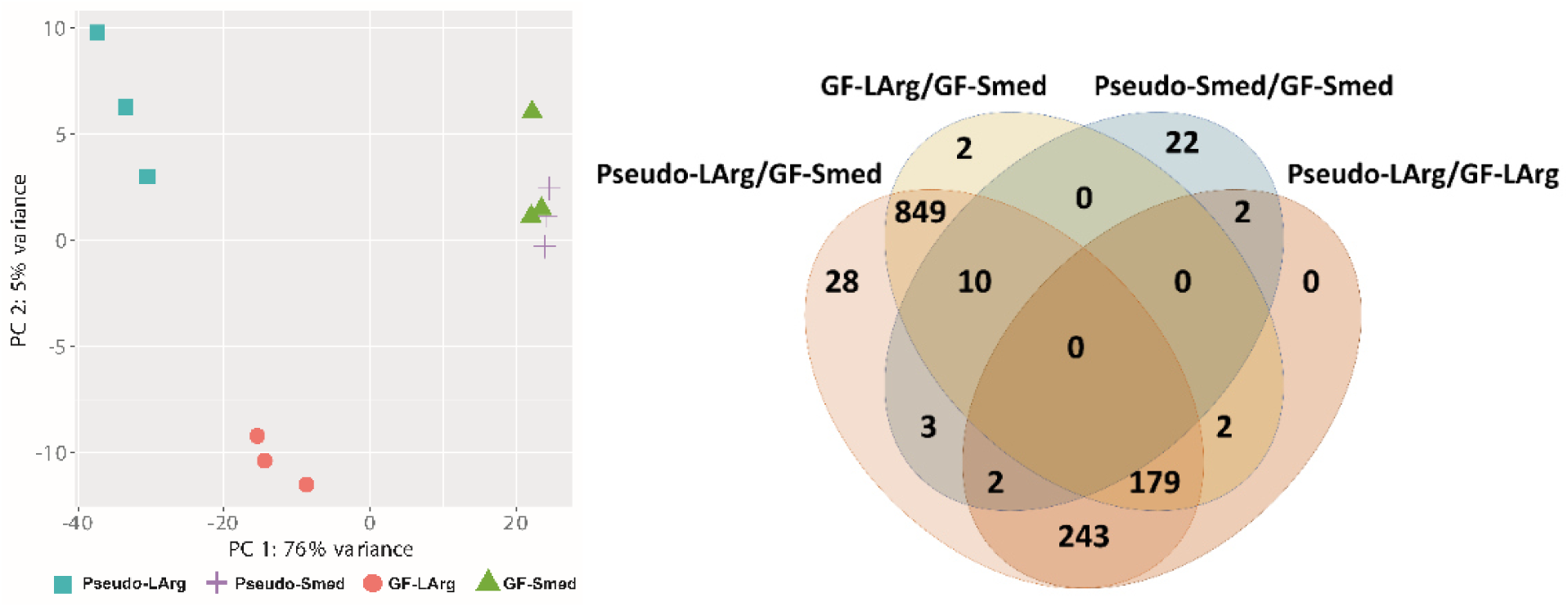
Transcriptional analysis of diseased *Hydra* 24 h post exposure. (Left) Principle component analysis (PCA) plot illustrating transcriptional differences between mono-colonised (*Pseudomonas alcaligenes* T3 (Pseudo)) and germ-free (GF) *Hydra* polyps exposed to L-arginine (LArg) or nutrient-deficient water (Smed). (Right) Venn-diagram illustrating differential regulation of annotated genes in different treatments.

**Fig. S13.**
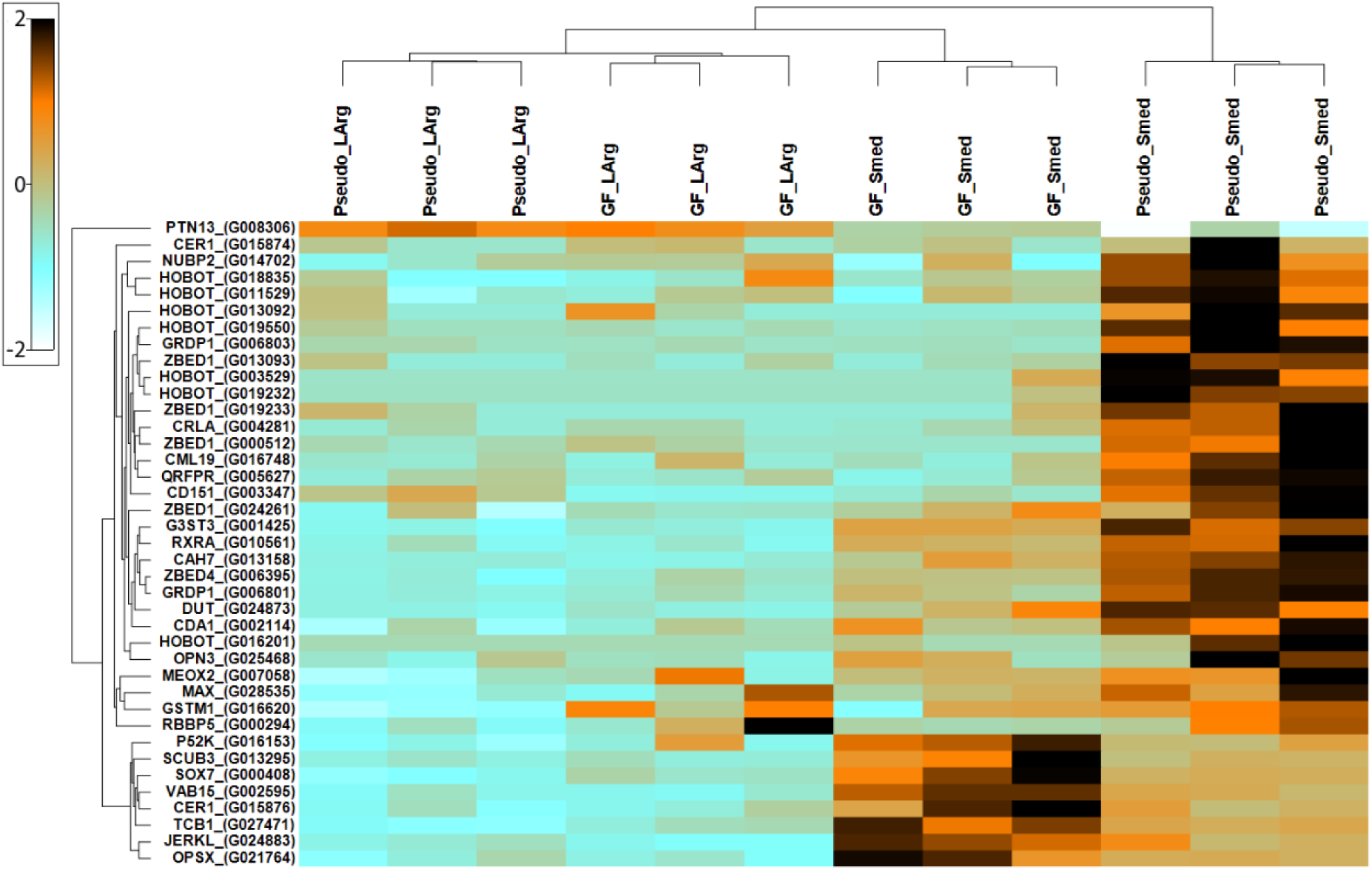
Heatmap of differential expressed all annotated genes that are differentially expressed (based on Z-score values) in *Hydra* polyps mono-colonised with *Pseudomonas alcaligenes* T3 (Pseudo) exposed to either L-arginine (LArg) or nutrient-deficient water (Smed). This was compared to germ-free (GF) polyps treated the same way.

**Fig. S14.**
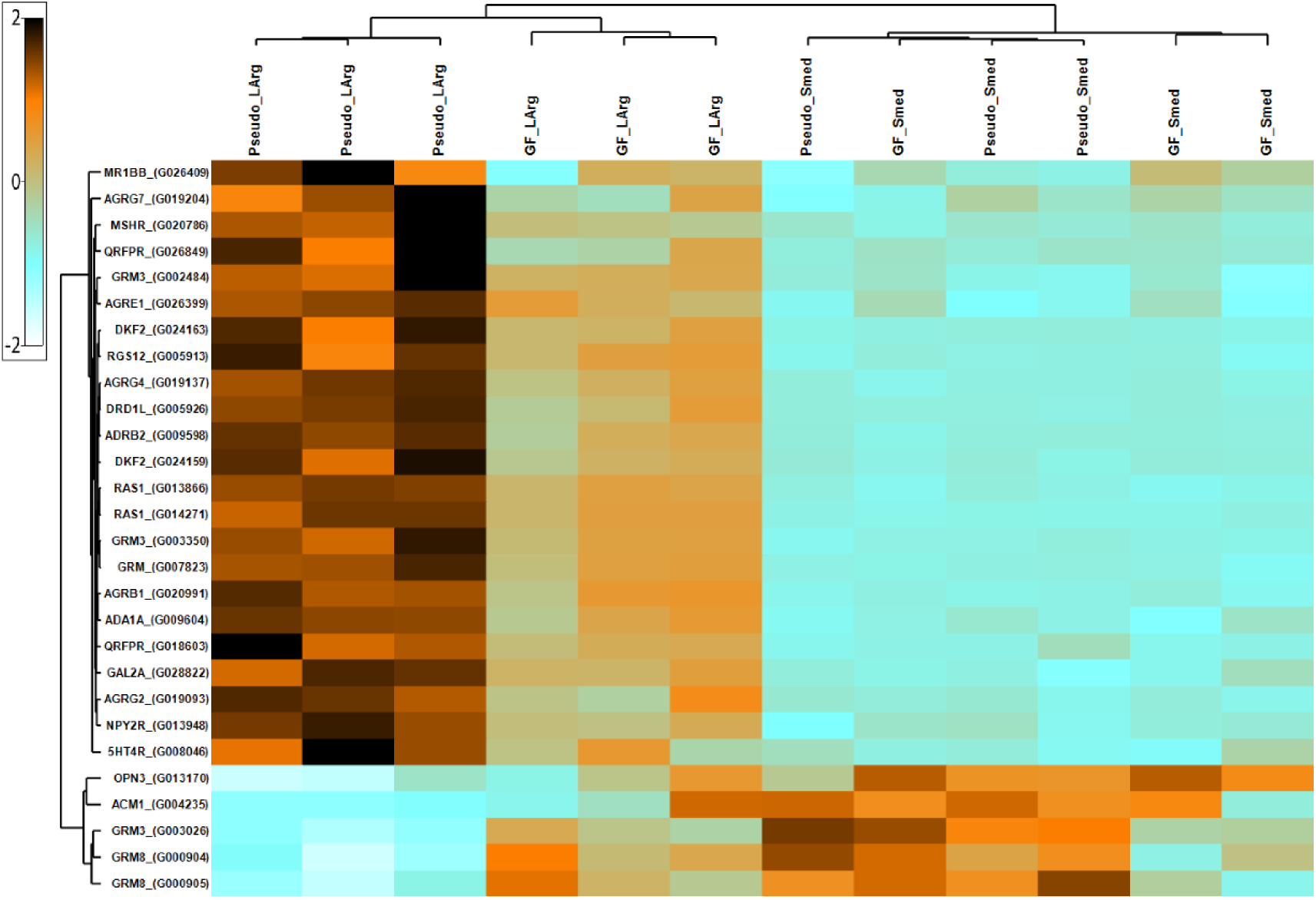
Heatmap of differential expressed G-proteins (based on Z-score values) in *Hydra* polyps mono-colonised with *Pseudomonas alcaligenes* T3 (Pseudo) exposed to either L-arginine (LArg) or nutrient-deficient water (Smed). This was compared to germ-free (GF) polyps treated the same way.

**Fig. S15.**
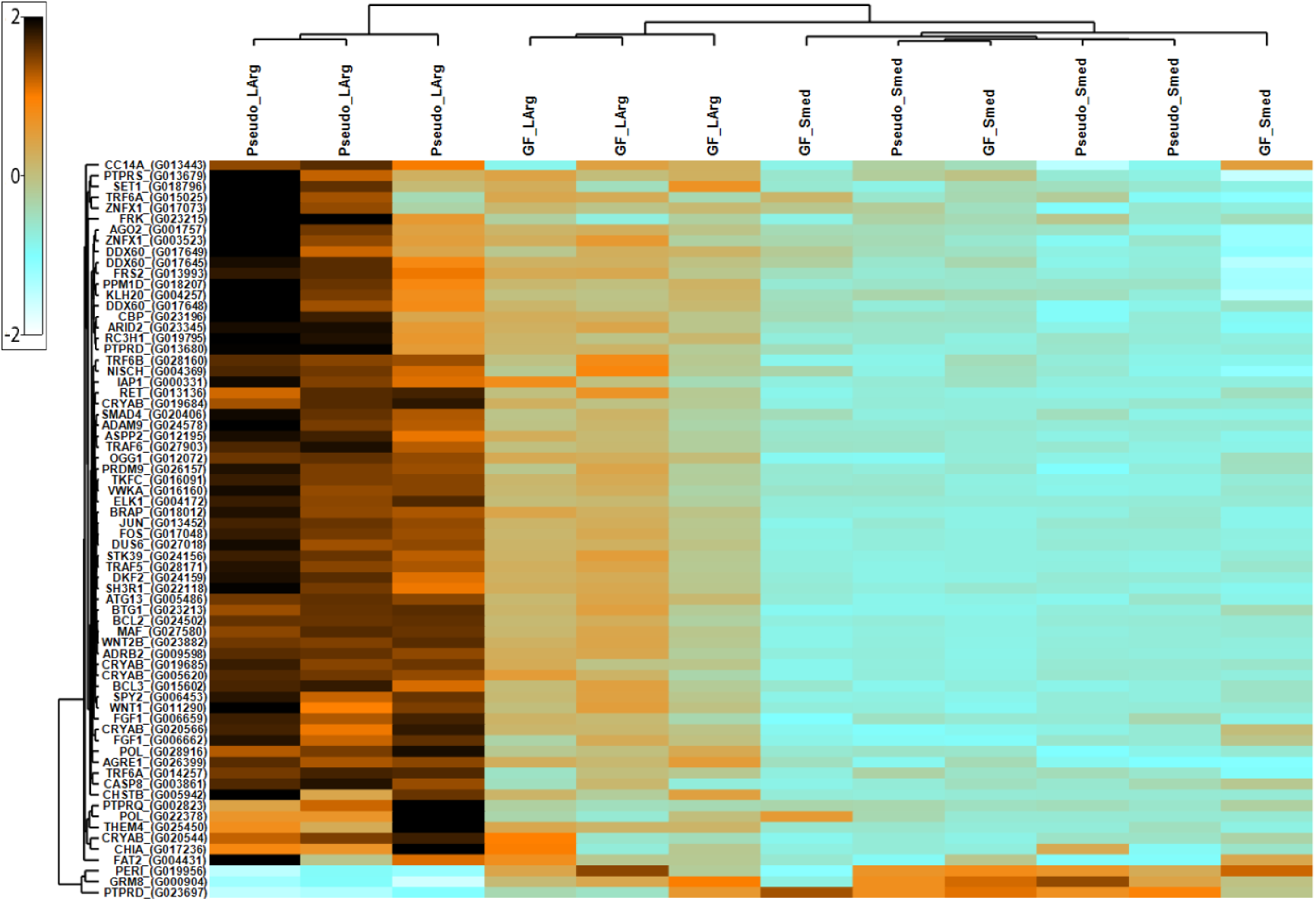
Heatmap of differential expressed immune responsive genes (based on Z-score values) in *Hydra* polyps mono-colonized with *Pseudomonas alcaligenes* T3 (Pseudo) exposed to either L-arginine (LArg) or nutrient-deficient water (Smed). This was compared to germ-free (GF) polyps treated the same way.

**Table S1.** Nutrient load in mg/l of sterile (0.02 µm) filtered lake water.

|  | **DOC** | **Phosphate** | **Nitrogen** | **C/N** |
| --- | --- | --- | --- | --- |
| Postsee | 9.1 | 0.044 | 1.10 | 8.3 |
| Lanker See | 8.9 | 0.014 | 0.78 | 11.4 |
| Tresdorfer<br>See | 10.0 | 0.011 | 2.00 | 5.0 |
| Selenter See | 6.4 | 0.005 | 0.57 | 11.2 |
| Plußsee | 6.9 | 0.017 | 0.89 | 7.8 |
| Schluensee | 0.1 | 0.006 | 0.46 | 0.2 |

